# The Labbing Project: A database management application for neuroimaging research

**DOI:** 10.1101/2023.01.02.522426

**Authors:** Zvi Baratz, Yaniv Assaf

## Abstract

The goal of this article is to present “The Labbing Project”; a novel neuroimaging data aggregation and preprocessing web application built with Django and VueJS. Neuroimaging data can be complex and time-consuming to work with, especially for researchers with limited programming experience. This web application aims to streamline the process of aggregating and preprocessing neuroimaging data by providing an intuitive, user-friendly interface that allows researchers to upload, organize, and preprocess their data with minimal programming requirements. The application utilizes Django, a popular Python web framework, to create a robust and scalable platform that can handle large volumes of data and accommodate the needs of a diverse user base. This robust infrastructure is complemented by a user-friendly VueJS frontend application, supporting commonplace data querying and extraction tasks. By automating common data processing tasks, this web application aims to save researchers time and resources, enabling them to focus on their research rather than data management.

## Introduction

Centralized data collection and distribution efforts are key to the advancement of neuroscientific research (Laird, 2021; Milham et al., 2018; Van Horn & Toga, 2009). Larger publicly available samples enable the detection and study of phenomena with smaller effect sizes more rapidly, by more people, and with more confidence (Laird, 2021; Madan, 2021; Smith & Nichols, 2018). Data sharing enhances the reproducibility of findings, reduces time and costs for new research, promotes transparency, and encourages collaboration across fields (Madan, 2017; Mar et al., 2013; Milham et al., 2018; Poldrack & Gorgolewski, 2014). In recent years, such public datasets have become more numerous and increasingly accessible (Borghi & Gulick, 2018; Horien et al., 2021; Laird, 2021).

This unprecedented scale and availability of data has a proven impact on the field of neuroimaging (Milham et al., 2018). However, there are still numerous issues hindering the ability of researchers to fully leverage these assets to optimize the quantity and quality of their scientific findings, individually or in collaboration (Borghi & Gulick, 2018; Routier et al., 2021). The management of subject information and both raw and packaged MRI data (i.e., converted to NIfTI and organized to conform with the BIDS standard) along with preprocessing results and any other, non-MRI data, in scale, can quickly prove to be a more complicated task than often anticipated. In one report, it is estimated that it generally takes two or three researchers working on a project based on data from public datasets roughly 6 to 9 months to download, process, and prepare the data for analysis (Horien et al., 2021). This period of time is also fertile ground for the introduction of more degrees of freedom to the analytical workflow (as well as human error), severely undermining the reproducibility of the results (Botvinik-Nezer et al., 2020; Maier-Hein et al., 2017). To better facilitate large-scale neuroimaging research, dedicated data and analysis management solutions are required.

This paper will explore the pipeline and methods meant to satisfy the following requirements:

1. General research information management
2. Raw DICOM metadata extraction and database management
3. Data conversion and standardization
4. Efficient querying and distribution functionality
5. Analysis workflow registration and orchestration

See the supplementary material (RDM Concept and Requirements) for a more detailed overview.

### Free and Open-source Software (FOSS) and Collaborative Science

The benefit of FOSS and collaborative software development on scientific research in general (Fortunato & Galassi, 2021) and neuroscientific research in particular (Gleeson et al., 2017; Halchenko & Hanke, 2012; White et al., 2019) has become a central topic of discussion in recent years. Working with and collaborating on open-source code repositories, including version control, testing, documentation, continuous integration (CI), and much more, is rapidly becoming a fundamental part of the technical capability expected from researchers in the field (Muller et al., 2015). Relevant educational content has become effectively ubiquitous in workshops and other educational programs, and an increasing number of contemporary initiatives are embracing the community-oriented philosophy and standards. This shift in the zeitgeist is largely owed to the widespread adoption of Python and the general-purpose programming language tools and paradigms it offers and encourages.

For the purposes of this paper, the desired RDM solution is one that is entirely free and open-source, and is designed as a collaborative, community-oriented software development project. Because most researchers are not experienced with commonplace web development technologies (e.g., NodeJS, PHP, Java, etc.), as much of the codebase should be written in Python as possible. This will encourage user involvement, foster awareness of infrastructural details and processing decisions, and ultimately serve to support the credibility and reproducibility of the derived findings.

### Existing Solutions

A number of neuroimaging RDM systems have been developed to support particular research centers and collaborations across the world. Some of the most notable examples are the Extensible Neuroimaging Archive Toolkit (XNAT, see https://www.xnat.org/) (Herrick et al., 2016; Marcus et al., 2007), Longitudinal Online Research and Imaging System (LORIS, see https://loris.ca/) (Das et al., 2012), and the newest addition, brainlife.io. All three are open-source and in active development. However, none offered both the required functionality and the means to create a sustainable, independent deployment of the application without specialized expertise at the time of writing. Table 1 offers a summary of the technological stacks and DICOM or BIDS-related functionality provided by these applications.

**Table 1:** A summary of existing open-source neuroimaging RDM applications’ technological stacks and the required DICOM or BIDS-related functionality. All information was independently gathered from openly accessible project websites, documentation, or code repositories.

| Name | Technological Stack | DICOM Import | DICOM to BIDS Conversion | BIDS Import | Deployment Documentation | Source Code |
| --- | --- | --- | --- | --- | --- | --- |
| XNAT | Linux<br>Traefik<sup>[1]</sup><br>PostgreSQL<sup>[2]</sup><br>Apache Tomcat<sup>[3]</sup> | ✓ | ✗ | ✗ | ✓ | bitbucket.org/xnatdev |
| LORIS | Linux<br>Apache HTTP Server<sup>[4]</sup><br>MySQL<sup>[5]</sup><br>PHP<sup>[6]</sup><br>React<sup>[7]</sup> | ✓ | ✗ | ✗ | ✓ | github.com/aces |
| brainlife.io | Mixed microservice architecture<br>Vue.js | ✗ | ✓* | ✓ | ✗ | github.com/brainlife |

\* brainlife.io has a new and exciting semi-automated web-based DICOM to BIDS conversion tool, *ezBIDS*, currently in development and approaching a stable release.
[1] https://traefik.io/ [2] https://www.postgresql.org/ [3] https://tomcat.apache.org/ [4] https://httpd.apache.org/ [5] https://www.mysql.com/ [6] https://php.net/ [7] https://reactjs.org/

The ability to manage a large DICOM dataset combined with external BIDS datasets and fluently associate the included or provided research information is a core requirement which is not yet properly handled by existing tools. In addition, none of the applications’ technological stacks are Python-based, making it significantly less likely for most researchers to be able to actively participate in their development. To overcome this technical difficulty, this article introduces a dedicated, Python-based, open-source and community-oriented RDM application.

## Materials and Methods

A dedicated [GitHub](https://www.github.com/) organization titled “The Labbing Project” (see https://github.com/TheLabbingProject) has been created to centralize development efforts. The following section will detail the technological stack used to build the application.

### Web Framework

Enabling secure remote access and providing robust infrastructure that can easily be deployed and scaled requires an uncompromising web application development framework. There are a number of Python-native high-level web frameworks (for a comprehensive review, see https://wiki.python.org/moin/WebFrameworks). However, the most popular option (by [GitHub](https://www.github.com/) star counts at the time of writing) is Django.

### Django

Django (see https://www.djangoproject.com/) is a widely used free and open-source web application development framework with several outstanding advantages. It is actively maintained by an independent non-profit organization and used by well-known websites, including [Instagram](https://www.instagram.com/), [Mozilla](https://www.mozilla.org/), [Pinterest](https://www.pinterest.com/), and more (see [Overview](https://www.djangoproject.com/start/overview/) page in the Django project’s website). The framework’s architecture promotes a modular design that is easily extensible and reusable. It also excels in database management, and includes a powerful Object-relational mapper (ORM) which greatly reduces the complexity of database administration and interaction by enabling the representation of data models as simple Python classes. Many aspects of routine web application development, such as web security, user administration, session management, caching, logging, testing, etc., are integrated into the framework in compliance with the highest standards in the field. The documentation site (see https://docs.djangoproject.com/) is exceptionally extensive and offers tutorials and educational content, thereby significantly reducing the learning-curve for potential contributors from the research community. Python’s renowned capacity for abstraction and intuitive syntax, combined with Django’s fully featured, scalable design, grants non-professional programmers unprecedented accessibility to actively participate in high-level web development.

Django web applications are written as a single *project*, providing general application-wide settings and resources, and *apps*, providing some custom, loosely coupled and reusable functionality (for more information, see the “[Applications](https://docs.djangoproject.com/en/dev/ref/applications/)” and “[Design philosophies](https://docs.djangoproject.com/en/dev/misc/design-philosophies/)” sections in the Django project’s documentation site). This architecture enables the creation of a particularly stable and maintainable foundation by tremendously facilitating the extension, modification, or replacement of different parts as requirements change over time or across deployments.

#### Application Programming Interface

Django REST framework (see https://www.django-rest-framework.org/) was used to generate a REST API for the application, exposing comprehensive database management and querying capabilities using simple web requests. In addition, returning generic, JSON-encoded responses enables flexible interaction with any number of external services or independent user interfaces.

### Application Server

An application server is used to translate HTTP requests between the web server (i.e., the machine running the web application) and the web application. *gunicorn* (see https://gunicorn.org/) is a free and open-source WSGI HTTP server commonly used for this purpose. It is fast, secure, highly configurable, and simple to use.

### Web Server

NGINX (see https://www.nginx.com/) is an open-source high-performance web server, reverse proxy, load balancer, and more. It is used as an intermediary between the web application and incoming requests and serves the returned static content. It also enables secure access over HTTPS using a TLS certificate.

### Database

Django officially supports numerous databases (see https://docs.djangoproject.com/en/dev/ref/databases/), with [PostgreSQL](https://www.postgresql.org/) being a leading choice for applications with higher demands in terms of database performance and flexibility.

### PostgreSQL

[PostgreSQL](https://www.postgresql.org/) is a free and open-source relational database management system (RDBMS). It is one of the most popular DBMSs in general (e.g., see https://db-engines.com/en/ranking), boasting some of the most advanced functionality in terms of both security and performance. Django’s [PostgreSQL](https://www.postgresql.org/) backend uses the psycopg (see https://www.psycopg.org/) Python adapter, which is written mostly in the C programming language, and provides a feature-rich, fast, and secure integration with the application’s components.

### Task Orchestration

Scalable execution of analytical workflows requires an advanced, distributed task queuing system. Preprocessing and feature extraction pipelines used in neuroimaging range from simple transformations to exceptionally resource-heavy, long-running tasks. Therefore, being able to initialize background processes across threads and machines in batch and monitor progress is an essential requirement.

### Celery

*[Celery](https://docs.celeryproject.org/)* is the most popular task queue manager written in Python (by [GitHub](https://www.github.com/) stars count at the time of writing). It is free and open-source, actively maintained, in widespread use in industry, and provides native support for integration with Django projects (see https://docs.celeryproject.org/en/stable/django/). By default, *[Celery](https://docs.celeryproject.org/)* uses RabbitMQ (see https://www.rabbitmq.com/) to send and receive messages, however, other message brokers such as Redis (see https://redis.io/) and Amazon SQS (see https://aws.amazon.com/sqs/) are also supported.

*django-celery-results* (see https://django-celery-results.readthedocs.io/) and *django-celery-beat* (see https://django-celery-beat.readthedocs.io/) are also enabled by default and complement *[Celery](https://docs.celeryproject.org/)* with database-integrated task monitoring and support for periodic task scheduling (respectively).

## Data Distribution

While the application’s REST API does provide endpoints for downloading both data instances and analysis results, distributing large collections of files necessitates a more controlled approach.

### Paramiko

*Paramiko* (see https://www.paramiko.org/) is a free and open-source Python implementation of the SSHv2 protocol. It enables highly controlled SFTP session negotiation over SSH, thereby allowing researchers to safely transport files to any SSH accessible machine.

### Front-end Framework

While Django does include a web template system (the Django Template Language, see https://docs.djangoproject.com/en/dev/topics/templates/) allowing for the generation of dynamic web pages, our attempts to leverage it and thereby keep the stack as purely Python-based as possible have failed to produce satisfactory results. In order for the application to offer a modern and responsive user experience, a dedicated front-end web development framework was deemed necessary.

### Vue.js

[Vue.js](https://vuejs.org/) is a popular free and open-source [JavaScript](https://www.javascript.com/) framework. It is lightweight, modular, rich in documentation and learning materials, and oriented towards both novice and expert developers. While some familiarity with HTML, CSS, and [JavaScript](https://www.javascript.com/) is required, [Vue.js](https://vuejs.org/) greatly simplifies the complexities of rendering web content in a browser. By interacting with the web application’s API, the front-end application offers a convenient graphical user interface (GUI) for researchers to query and retrieve data and derivatives from the server.

## Results

The following section will detail the components of the Django and [Vue.js](https://vuejs.org/) web application, available under the dedicated [GitHub](https://www.github.com/) organization’s page at https://www.github.com/TheLabbingProject.

### General Research Management Application

Even though it is built to support neuroimaging-based academic research, the core of the application is entirely domain-agnostic. Elementary research entities, such as subjects, studies, and experimental procedures, are managed independently of any particular data model.

#### pylabber

*pylabber* (see https://www.github.com/TheLabbingProject/pylabber) is the base project repository (see Django section in Materials and Methods) for the complete web application. In addition to centralizing configurations and resources, it includes two general-purpose research apps:

- *accounts:* Manages laboratory and researcher information, as well as data distribution. Administrators can register new laboratories and users (researchers), and users can provide SSH credentials (used to negotiate SFTP transport sessions) to remote machines, termed “export destinations”. Raw data as well as analysis results export methods are provided, and *[Celery](https://docs.celeryproject.org/)* tasks are available for initiating controlled and monitored export sessions as background processes.
- *research:* Manages general research-related entities. Relies on external apps to associate models representing data acquisitions (such as an MRI session) to the appropriate subject, and models representing data instances (such as an MRI scan) to study groups. The decision to move group association from subjects to data instances was made to allow for flexible association of any part of the acquired data to any number of study groups. Each study may be assigned a number of study groups, as well as a number of experimental procedures. An experimental procedure consists of an ordered sequence of steps which may be categorized as either tasks or data acquisitions (see the *research* container nested within the *pylabber* container and highlighted in purple in Fig. 1).

**Fig. 1:**
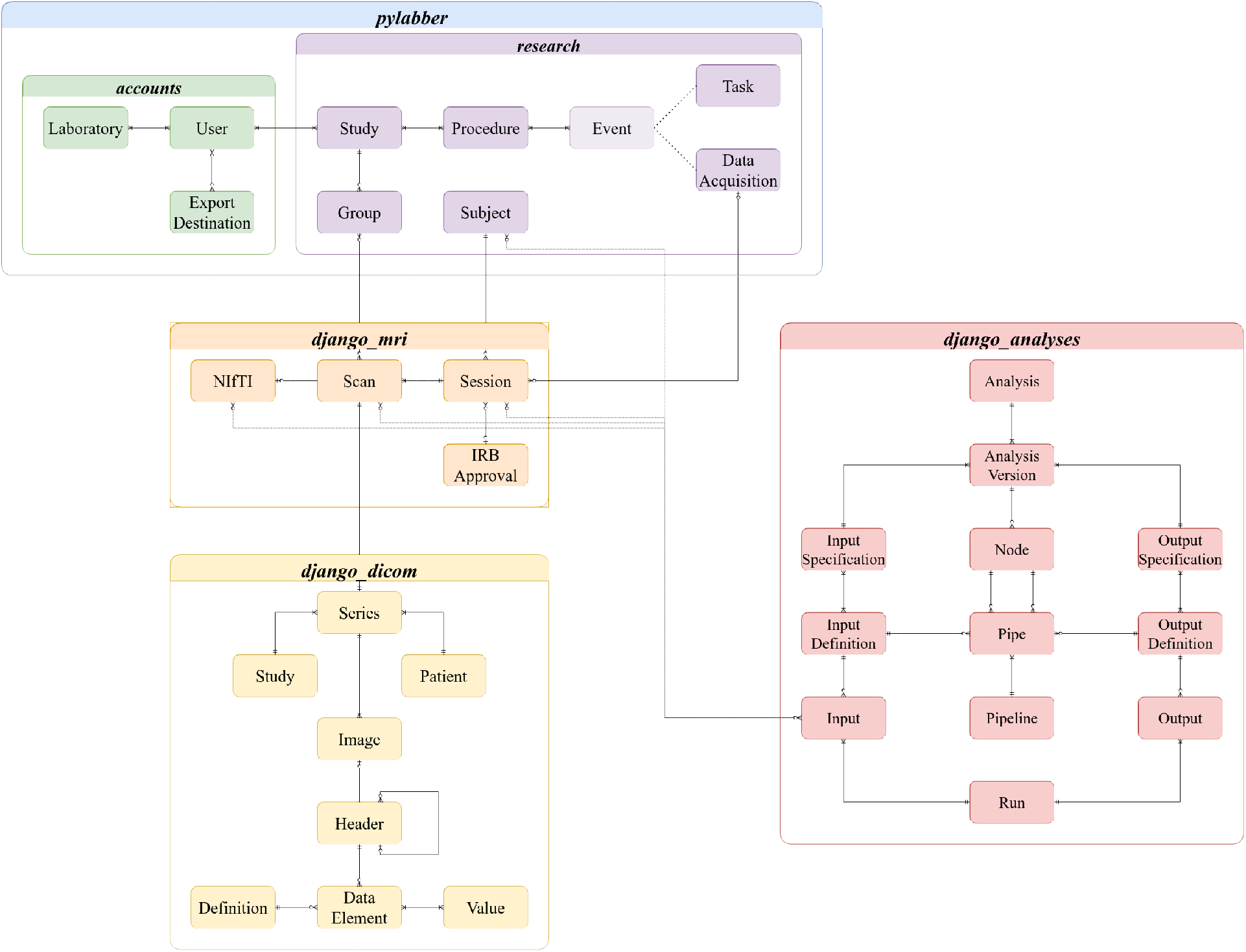
Crow’s foot Entity-Relationship (ER) diagram illustrating the database tables managed by the application and relationships between them.

##### Admin Interface

Django provides powerful automated admin interface generation utilities (see https://docs.djangoproject.com/en/dev/ref/contrib/admin/). The application makes extensive use of these features to provide highly detailed admin access to all the application’s models (i.e., database tables) including CRUD operations, useful filters, and summary plots.

**Fig. 2:**
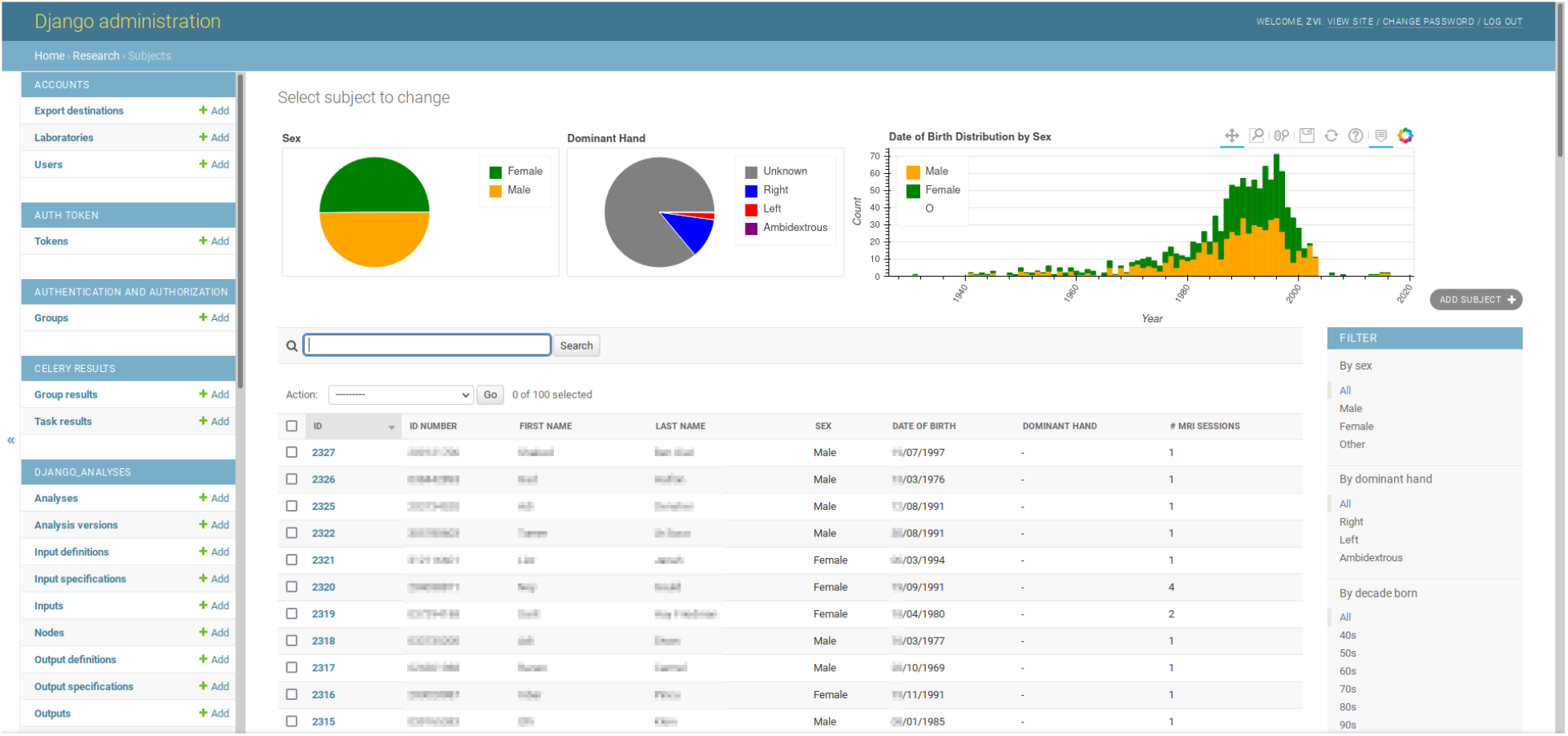
The subject model’s admin interface list view. Sex, dominant hand, and date of birth distribution plots are displayed to offer a visual summary of the displayed queryset. A general search field allows searching by first name, last name, or ID number. More filters are available on the right panel. Other models’ list views are available on the left panel.

### Reusable Django Apps

The integration of references to files and database access to metadata information is handled using reusable Django apps and conforms with the Django framework’s general design philosophy. Other than reusability, encapsulation of domain-specific logic also contributes to both the maintainability and extensibility of the code-base.

#### django_analyses

*django_analyses* provides database-driven analysis registration and execution management (see the *django_analyses* container, highlighted in red in Fig. 1).

##### Analysis Integration

Analysis interfaces (i.e., Python classes or functions used to initialize the execution of an analysis) are registered with a particular analysis version, which represents it in the database and holds references to the appropriate input and output specifications. When an analysis version is executed, a new run instance is created to combine the information about the analysis version with the provided inputs and eventually outputs (or traceback information if an exception was raised). If the same analysis version with the same input configuration is required again, the existing run instance is returned.

Node instances are used as execution templates and provide a constant reference to runs of a specific analysis version with a particular configuration. Controlled execution of analyses with *[Celery](https://docs.celeryproject.org/)* is handled by providing nodes with input data and returning the created (or existing) run instances.

##### Pipeline Integration

Pipelines, or workflows, may be registered in the database as a sequence of pipes. Each pipe describes the flow of data from one node’s output to another node’s input specification. A dedicated “pipeline runner” class is provided by the app to control the ordered execution of nodes according to the dependencies outlined by the pipes.

For more information about *django_analyses* and complete usage examples, see the app’s documentation site (available at https://django-analyses.readthedocs.io/).

#### django_dicom

*django_dicom* is used to import *.dcm* files, extract metadata from the headers, and keep an organized archive of the included DICOM entities (see [Part 3 Section A.1.2](https://dicom.nema.org/dicom/2013/output/chtml/part03/chapter_A.html) of the DICOM Standard as well as the *django_dicom* container, highlighted in yellow in Fig. 1). All attributes associated with each information entity included in a DICOM header are serialized to the database together with principal acquisition parameters (e.g., TE, TI, TR, etc.).

#### dicom_parser

Header information is read using *dicom_parser* (see https://www.github.com/open-dicom/dicom_parser), a wrapper around *pydicom* (see https://pydicom.github.io/) developed specifically for the purposes of the app. While *pydicom* focuses on low-level header parsing in compliance with the DICOM Standard, *dicom_parser* provides a higher-level API to facilitate type conversion and vendor-specific metadata extraction methods. In addition, *dicom_parser* includes a “sequence detection” utility, enabling the configuration of commonplace heuristics for automated categorization of the scanning protocol (e.g., MPRAGE, FLAIR, fMRI, dMRI, etc.).

#### django_mri

*django_mri* provides an abstraction layer which is ultimately meant to keep the users practically agnostic of file-format and metadata extraction processes (see the *django_mri* container, highlighted in orange in Fig. 1). It uses *django_dicom* alongside a supplementary NIfTI database table to simplify interaction with MRI data by providing abstract “scan” and “session” representations. The scan data model keeps a reference to source files and automates whatever possible to allow shifting the focus from infrastructure and technical details to processing and results.

In terms of file management, *dicom_parser*’s sequence detection functionality provides information which is essential for automated BIDS-compliant organization. By taking advantage of the configured heuristics, *django_mri* maintains a “global” BIDS dataset, which holds the combined datasets of multiple studies. All research information is maintained in the database and any available BIDS app may be applied (see https://bids-apps.neuroimaging.io/).

Finally, *django_mri* makes extensive use of *django_analyses* and *[nipype](https://nipype.readthedocs.io/)* (Gorgolewski et al., 2011) to provide the specifications and analysis interfaces (see Analysis Integration) for numerous neuroimaging processing tools, including *[fMRIPrep](https://fmriprep.org/)* (Esteban et al., 2019), *FreeSurfer* (Fischl, 2012), *[QSIPrep](https://qsiprep.readthedocs.io/)* (Cieslak et al., 2021), *CAT* (see http://www.neuro.uni-jena.de/cat/), and more.

### Front-end Application

#### vuelabber

A front-end application created with [Vue.js](https://vuejs.org/) is used to provide users with a modern-looking and highly functional graphical user interface. Communication between the front-end and the server is based on the application’s REST API (see Application Programming Interface) and any part of it may easily be customized, extended, or replaced.

**Fig. 3:**
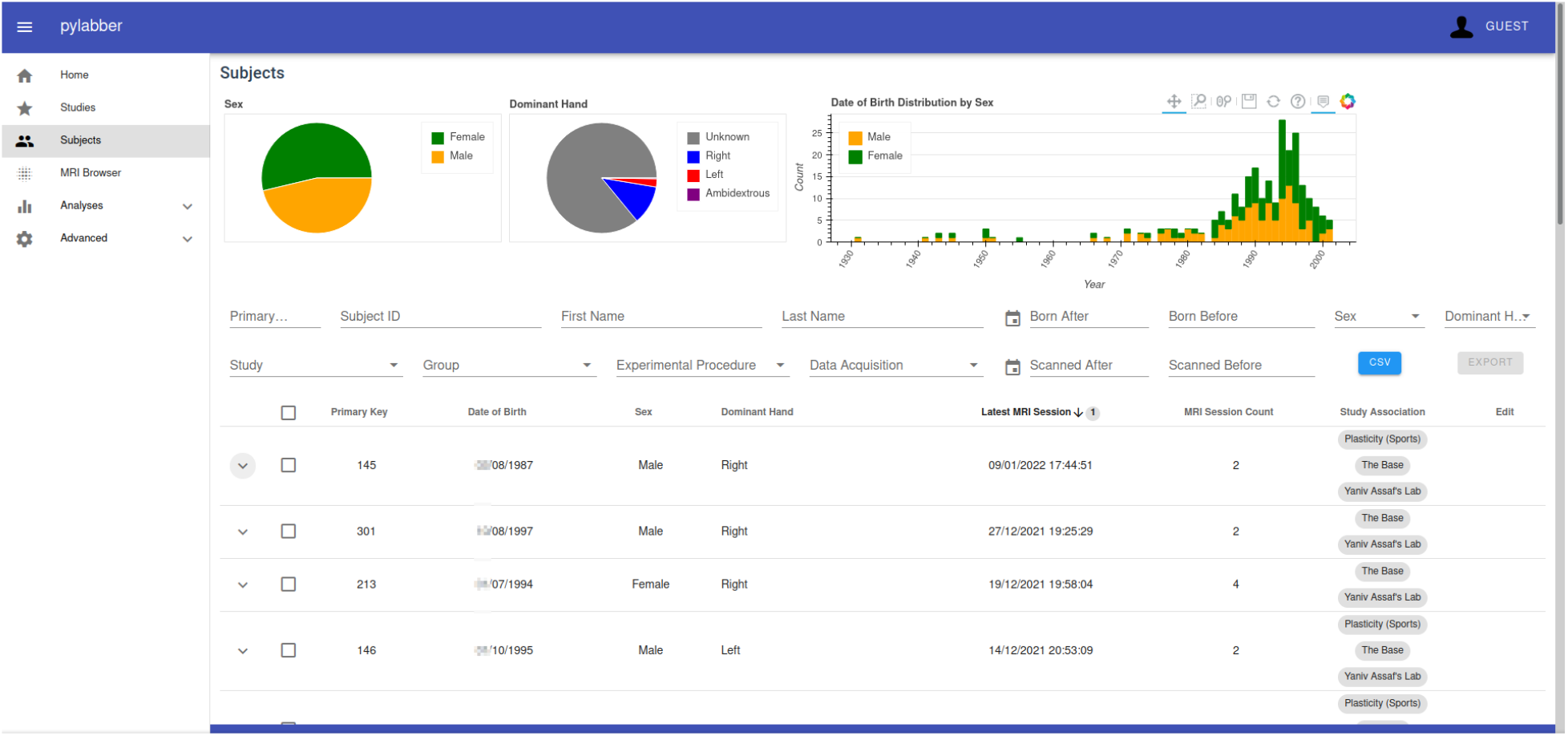
Subjects list view as displayed in the [Vue.js](https://vuejs.org/) front-end application. More detailed research information and filters are available compared to the admin interface, along with the same summary plots.

### Neuroimaging Research Database

The purpose of the application is to manage a large and highly variable MRI dataset alongside relevant research information and derivatives. Currently, a single, local deployment (self-managed on a physical server) of the application is used to manage all independently resourced data, as well as other shared resources (e.g., the HCP dataset) in our laboratory. This includes over 4000 MRI scanning sessions from over 3000 subjects (including the HCP dataset), as well as over 7000 preprocessing pipeline runs.

## Discussion

The field of neuroscience is in the process of adapting to a sharp transition in the scale of available data. To sustain this transition, the research community must not only overcome technical and statistical difficulties, but also challenges of communication and collaboration.

Centralized management, analysis, and distribution of data is rapidly becoming the foundation of the great majority of imaging studies, and this trend is reasonably expected to become effectively ubiquitous is the foreseeable future. A number of research data management (RDM) solutions with the purpose of facilitating large-scale neuroimaging-based research have already emerged, and are making a substantial difference in the average sample sizes reported in published literature. While curating data and making it available for researchers remains the obvious prerequisite for progress, it will be the advancement of infrastructural tools enabling the distribution and aggregation of data in scale that will determine the slope.

This article introduces a novel open-source and Python-based neuroimaging RDM application, termed “The Labbing Project”. What makes this particular software suite distinct is that it is written first and foremost as a community-oriented collection of modular tools, meant to be maintained and developed by the academic audience itself. The purpose of this project is not to provide researchers with a definite solution to all research needs, rather, it is meant to encourage involvement and standardization of archiving, preprocessing, and data distribution methodologies, thereby facilitating and optimizing collaboration efforts. In other words, “The Labbing Project” is an ongoing effort to abstractify common research entities and processes, as defined and utilized by researchers, to enable the exploration of larger datasets with higher reliability.

Other than choosing the Python as the core programming language of the application, extensive efforts have been made to make the codebase as accessible to researchers as possible. All included packages have detailed documentation sites, including installation instructions, basic usage, and in depth information regarding their respective structure and API. In accordance with Python’s general philosophy, Django’s object-oriented and modular design greatly simplifies many aspects of web application frameworks, significantly reducing the associated learning curve. In addition, there is a strong emphasis and compliance with industry-level coding standards and consistently striving for highly readable and comprehensively tested code.

As disciplinary conventions grow and evolve, researchers are increasingly expected to become proficient in various aspects of software development and database management. These skills are often taught in seminars and workshops, rather than in any professional environment, and fittingly, the great majority of research findings are based on code that would not be considered “production-ready” in an industrial setting. To create a foundation for sustainable progress, researchers must collaborate and participate in the development of a transparent and uncompromising technological stack. “The Labbing Project” is an example of one such proposed stack and the components it consists of.

## Supplementary Material

### RDM Concept and Requirements

A complete neuroimaging research data management (RDM) solution would be one that enables researchers to seamlessly integrate raw, independently acquired data, processing workflows, and results, with publicly shared datasets and their corresponding derivatives. Generally, a neuroimaging data acquisition is assumed to be an MRI session originally encoded in the DICOM format, but most commonly converted to NIfTI and packaged as part of a BIDS-compliant dataset. The fundamental requirements from an RDM system would include:

1. General research information management. Collected data needs to be associated with a particular study group (e.g., general population, control, task, etc.), and studies should be assigned collaborators (researchers), thereby granting access. The entities and relationships required to model the research process adequately may be conceptualized and illustrated in a great number of ways, Fig. 4 proposes one such conceptual model.
2. Raw DICOM metadata extraction and database management. Independently resourced data is assumed to be encoded in the DICOM format, and therefore raw data files include a header section containing study and subject-relevant information. The ability to import *.dcm* files and maintain a database of the associated DICOM entities (see [Part 3 Section A.1.2](https://dicom.nema.org/dicom/2013/output/chtml/part03/chapter_A.html) of the DICOM Standard) is core to the workflow of the great majority of neuroimaging laboratories. Even after metadata extraction and conversion, DICOM files must be archived and made efficiently queryable for future inspection. Other than the general best-practice of never deleting raw data, information about the influence of acquisition parameters on derived measurements is constantly being updated (McNabb et al., 2020; Risk et al., 2018; Todd et al., 2016), along with metadata sharing standards and even conversion methods.
3. Data conversion and standardization. As established, raw DICOM data aggregated in *.dcm* files needs to be converted to NIfTI and packaged to comply with the BIDS standard. *dcm2niix* (Li et al., 2016) (https://github.com/rordenlab/dcm2niix) provides exceptionally reliable format-conversion functionality, including JSON-sidecar generation (as specified in the BIDS standard). Once converted, the generated *.nii* and *json* files must remain indefinitely associated with their original representation as a DICOM series. Organization of the files as a BIDS-compliant dataset, however, is still only partially automated. While several tools offering semi-automated configurable workflows (e.g., *HeuDiConv, Dcm2Bids*) or interactive GUIs (see *ezbids*, available at https://brainlife.io/ezbids/ (Avesani et al., 2019) are in continuous development, a completely automated solution has yet to emerge. In addition, the RDM would need to be able to reorganize and validate the standardization of data “pulled” from external repositories. For BIDS-compliant datasets, this can be handled relatively easily by serialization of the encoded metadata to the database alongside minor reorganization or renaming of the source directories. Other cases, however, such as the HCP, would require more customized translation methods.
4. Efficient querying and distribution functionality. Once archived and packaged, the data must become accessible to the appropriate researchers. The application needs to provide comprehensive querying capabilities, covering both research entity relations and raw acquisition metadata. In addition, once a queryset has been curated, some method of distribution is required to export the data dependably to remote servers or workstations.
5. Analysis workflow registration and orchestration. While not strictly in the scope of an RDM, collaboration on a large neuroimaging dataset mandates a shared analytical workflow management tool. The scale of the data and resources required to preprocess it render the raw dataset practically unusable to most researchers (Plis et al., 2016). Instead, administrators should be able to register analytical pipelines, execute them, and enable access to the results. In addition to enabling researchers to study a much larger dataset than otherwise possible and the more economical allocation of resources, as well as the significant enhancement in reproducibility, having all workflows and their parameters documented and organized provides an opportunity to better understand the effects of various preprocessing parameters on derived results.

**Fig. 4:**
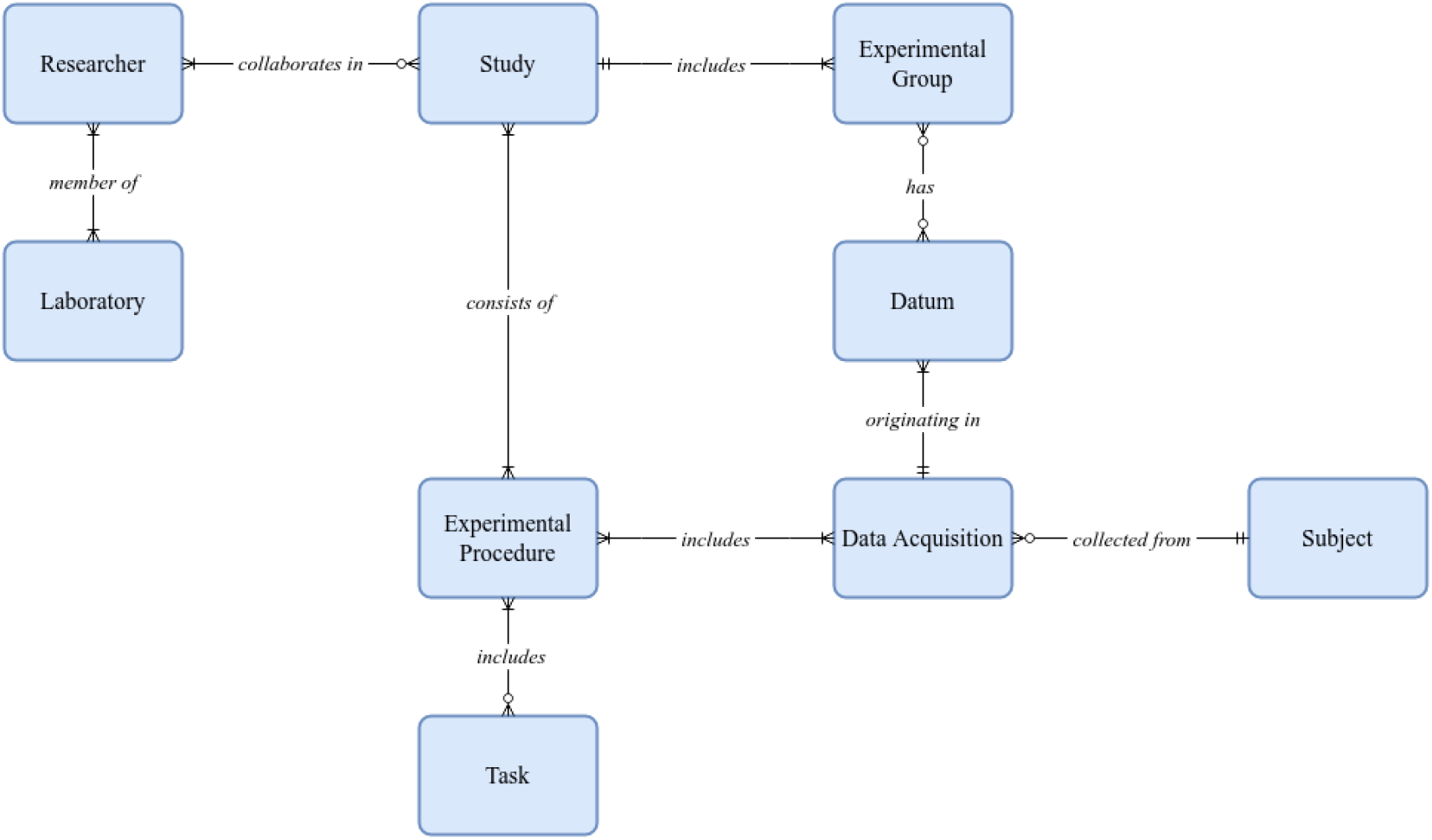
Proposed conceptual crow’s foot Entity-Relationship (ER) diagram illustrating research data management entities and their relationships.

**Fig. 5:**
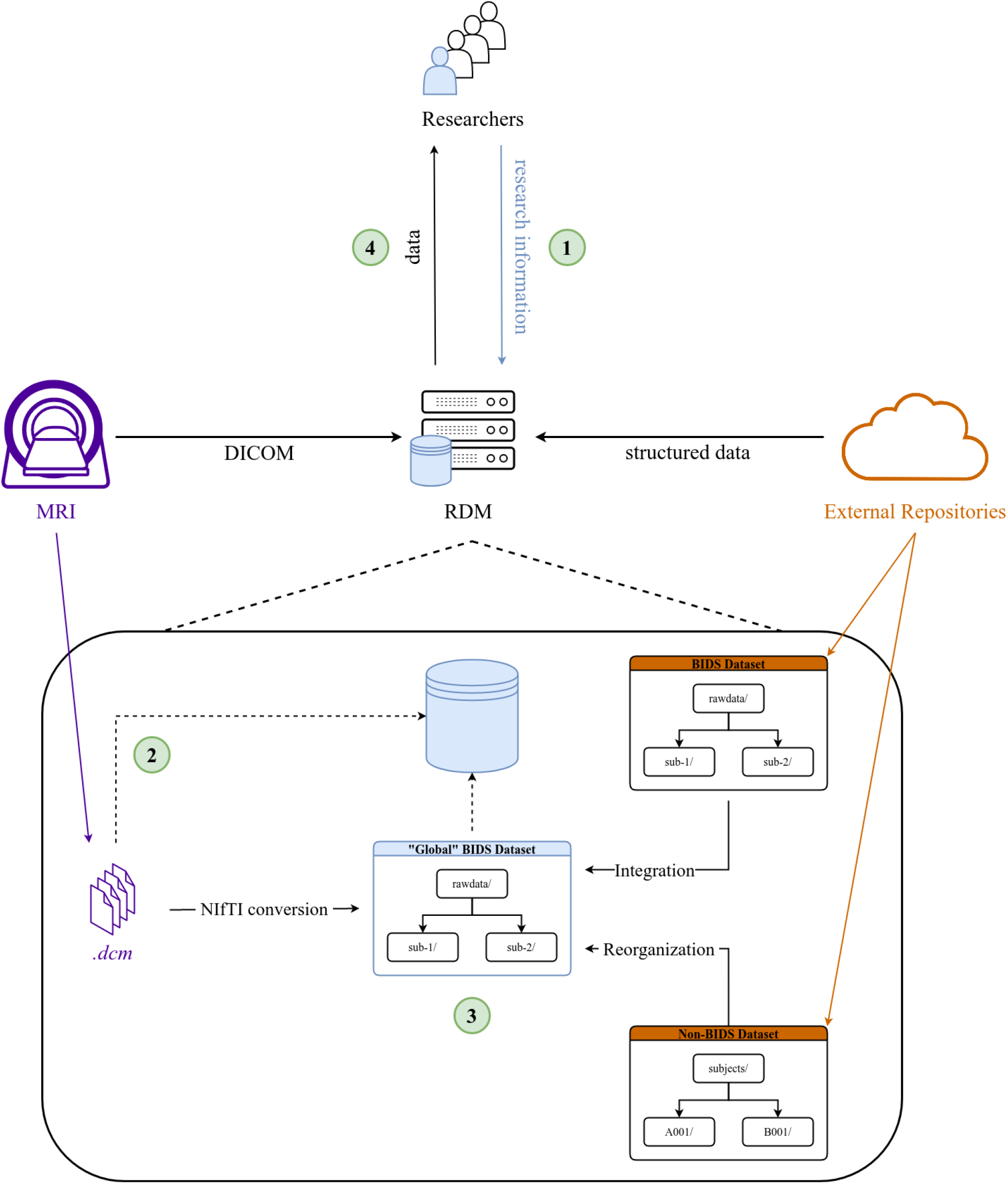
Schematic representation of the fundamental requirements from a neuroimaging RDM system. Numbers circled in green correspond to the numbers of the requirements as listed above. Blue user represents an admin or laboratory manager providing general research information.

## Notes

https://github.com/TheLabbingProject

### Competing Interest Statement

The authors have declared no competing interest.

### Summary of Updates

Removed duplicate paragraph.

## Bibliography

Avesani, P., McPherson, B., Hayashi, S., Caiafa, C. F., Henschel, R., Garyfallidis, E., Kitchell, L., Bullock, D., Patterson, A., Olivetti, E., Sporns, O., Saykin, A. J., Wang, L., Dinov, I., Hancock, D., Caron, B., Qian, Y., & Pestilli, F. (2019). The open diffusion data derivatives, brain data upcycling via integrated publishing of derivatives and reproducible open cloud services. Scientific Data, 6(1), 69. https://doi.org/10.1038/s41597-019-0073-y

Borghi, J. A., & Gulick, A. E. V. (2018). Data management and sharing in neuroimaging: Practices and perceptions of MRI researchers. PLOS ONE, 13(7), e0200562. https://doi.org/10.1371/journal.pone.0200562

Botvinik-Nezer, R., Holzmeister, F., Camerer, C. F., Dreber, A., Huber, J., Johannesson, M., Kirchler, M., Iwanir, R., Mumford, J. A., Adcock, R. A., Avesani, P., Baczkowski, B. M., Bajracharya, A., Bakst, L., Ball, S., Barilari, M., Bault, N., Beaton, D., Beitner, J., … Schonberg, T. (2020). Variability in the analysis of a single neuroimaging dataset by many teams. Nature, 582(7810), 84–88. https://doi.org/10.1038/s41586-020-2314-9

Cieslak, M., Cook, P. A., He, X., Yeh, F.-C., Dhollander, T., Adebimpe, A., Aguirre, G. K., Bassett, D. S., Betzel, R. F., Bourque, J., Cabral, L. M., Davatzikos, C., Detre, J. A., Earl, E., Elliott, M. A., Fadnavis, S., Fair, D. A., Foran, W., Fotiadis, P., … Satterthwaite, T. D. (2021). QSIPrep: An integrative platform for preprocessing and reconstructing diffusion MRI data. Nature Methods, 18(7), Article 7. https://doi.org/10.1038/s41592-021-01185-5

Das, S., Zijdenbos, A., Vins, D., Harlap, J., & Evans, A. (2012). LORIS: A web-based data management system for multi-center studies. Frontiers in Neuroinformatics, 5. https://www.frontiersin.org/article/10.3389/fninf.2011.00037

Esteban, O., Markiewicz, C. J., Blair, R. W., Moodie, C. A., Isik, A. I., Erramuzpe, A., Kent, J. D., Goncalves, M., DuPre, E., Snyder, M., Oya, H., Ghosh, S. S., Wright, J., Durnez, J., Poldrack, R. A., & Gorgolewski, K. J. (2019). fMRIPrep: A robust preprocessing pipeline for functional MRI. Nature Methods, 16(1), Article 1. https://doi.org/10.1038/s41592-018-0235-4

Fischl, B. (2012). FreeSurfer. NeuroImage, 62(2), 774–781. https://doi.org/10.1016/j.neuroimage.2012.01.021

Fortunato, L., & Galassi, M. (2021). The case for free and open source software in research and scholarship. Philosophical Transactions of the Royal Society A: Mathematical, Physical and Engineering Sciences, 379(2197), 20200079. https://doi.org/10.1098/rsta.2020.0079

Gleeson, P., Davison, A. P., Silver, R. A., & Ascoli, G. A. (2017). A Commitment to Open Source in Neuroscience. Neuron, 96(5), 964–965. https://doi.org/10.1016/j.neuron.2017.10.013

Gorgolewski, K., Burns, C. D., Madison, C., Clark, D., Halchenko, Y. O., Waskom, M. L., & Ghosh, S. S. (2011). Nipype: A Flexible, Lightweight and Extensible Neuroimaging Data Processing Framework in Python. Frontiers in Neuroinformatics, 5. https://doi.org/10.3389/fninf.2011.00013

Halchenko, Y., & Hanke, M. (2012). Open is Not Enough. Let’s Take the Next Step: An Integrated, Community-Driven Computing Platform for Neuroscience. Frontiers in Neuroinformatics, 6. https://www.frontiersin.org/article/10.3389/fninf.2012.00022

Herrick, R., Horton, W., Olsen, T., McKay, M., Archie, K. A., & Marcus, D. S. (2016). XNAT Central: Open sourcing imaging research data. NeuroImage, 124, 1093–1096. https://doi.org/10.1016/j.neuroimage.2015.06.076

Horien, C., Noble, S., Greene, A. S., Lee, K., Barron, D. S., Gao, S., O’Connor, D., Salehi, M., Dadashkarimi, J., Shen, X., Lake, E. M. R., Constable, R. T., & Scheinost, D. (2021). A hitchhiker’s guide to working with large, open-source neuroimaging datasets. Nature Human Behaviour, 5(2), 185–193. https://doi.org/10.1038/s41562-020-01005-4

Laird, A. R. (2021). Large, open datasets for human connectomics research: Considerations for reproducible and responsible data use. NeuroImage, 244, 118579. https://doi.org/10.1016/j.neuroimage.2021.118579

Li, X., Morgan, P. S., Ashburner, J., Smith, J., & Rorden, C. (2016). The first step for neuroimaging data analysis: DICOM to NIfTI conversion. Journal of Neuroscience Methods, 264, 47–56. https://doi.org/10.1016/j.jneumeth.2016.03.001

Madan, C. R. (2017). Advances in Studying Brain Morphology: The Benefits of Open-Access Data. Frontiers in Human Neuroscience, 11. https://www.frontiersin.org/article/10.3389/fnhum.2017.00405

Madan, C. R. (2021). Scan Once, Analyse Many: Using Large Open-Access Neuroimaging Datasets to Understand the Brain. Neuroinformatics. https://doi.org/10.1007/s12021-021-09519-6

Maier-Hein, K. H., Neher, P. F., Houde, J.-C., Côté, M.-A., Garyfallidis, E., Zhong, J., Chamberland, M., Yeh, F.-C., Lin, Y.-C., Ji, Q., Reddick, W. E., Glass, J. O., Chen, D. Q., Feng, Y., Gao, C., Wu, Y., Ma, J., He, R., Li, Q., … Descoteaux, M. (2017). The challenge of mapping the human connectome based on diffusion tractography. Nature Communications, 8(1), Article 1. https://doi.org/10.1038/s41467-017-01285-x

Mar, R. A., Spreng, R. N., & DeYoung, C. G. (2013). How to produce personality neuroscience research with high statistical power and low additional cost. Cognitive, Affective, & Behavioral Neuroscience, 13(3), 674–685. https://doi.org/10.3758/s13415-013-0202-6

Marcus, D. S., Olsen, T. R., Ramaratnam, M., & Buckner, R. L. (2007). The extensible neuroimaging archive toolkit. Neuroinformatics, 5(1), 11–33. https://doi.org/10.1385/NI:5:1:11

McNabb, C. B., Lindner, M., Shen, S., Burgess, L. G., Murayama, K., & Johnstone, T. (2020). Inter-slice leakage and intra-slice aliasing in simultaneous multi-slice echo-planar images. Brain Structure and Function, 225(3), 1153–1158. https://doi.org/10.1007/s00429-020-02053-2

Milham, M. P., Craddock, R. C., Son, J. J., Fleischmann, M., Clucas, J., Xu, H., Koo, B., Krishnakumar, A., Biswal, B. B., Castellanos, F. X., Colcombe, S., Di Martino, A., Zuo, X.-N., & Klein, A. (2018). Assessment of the impact of shared brain imaging data on the scientific literature. Nature Communications, 9(1), Article 1. https://doi.org/10.1038/s41467-018-04976-1

Muller, E., Bednar, J. A., Diesmann, M., Gewaltig, M.-O., Hines, M., & Davison, A. P. (2015). Python in neuroscience. Frontiers in Neuroinformatics, 9. https://www.frontiersin.org/article/10.3389/fninf.2015.00011

Plis, S. M., Sarwate, A. D., Wood, D., Dieringer, C., Landis, D., Reed, C., Panta, S. R., Turner, J. A., Shoemaker, J. M., Carter, K. W., Thompson, P., Hutchison, K., & Calhoun, V. D. (2016). COINSTAC: A Privacy Enabled Model and Prototype for Leveraging and Processing Decentralized Brain Imaging Data. Frontiers in Neuroscience, 10. https://www.frontiersin.org/article/10.3389/fnins.2016.00365

Poldrack, R. A., & Gorgolewski, K. J. (2014). Making big data open: Data sharing in neuroimaging. Nature Neuroscience, 17(11), 1510–1517. https://doi.org/10.1038/nn.3818

Risk, B. B., Kociuba, M. C., & Rowe, D. B. (2018). Impacts of simultaneous multislice acquisition on sensitivity and specificity in fMRI. NeuroImage, 172, 538–553. https://doi.org/10.1016/j.neuroimage.2018.01.078

Routier, A., Burgos, N., Díaz, M., Bacci, M., Bottani, S., El-Rifai, O., Fontanella, S., Gori, P., Guillon, J., Guyot, A., Hassanaly, R., Jacquemont, T., Lu, P., Marcoux, A., Moreau, T., Samper-González, J., Teichmann, M., Thibeau-Sutre, E., Vaillant, G., … Colliot, O. (2021). Clinica: An Open-Source Software Platform for Reproducible Clinical Neuroscience Studies. Frontiers in Neuroinformatics, 15, 689675. https://doi.org/10.3389/fninf.2021.689675

Smith, S. M., & Nichols, T. E. (2018). Statistical Challenges in “Big Data” Human Neuroimaging. Neuron, 97(2), 263–268. https://doi.org/10.1016/j.neuron.2017.12.018

Todd, N., Moeller, S., Auerbach, E. J., Yacoub, E., Flandin, G., & Weiskopf, N. (2016). Evaluation of 2D multiband EPI imaging for high-resolution, whole-brain, task-based fMRI studies at 3T: Sensitivity and slice leakage artifacts. NeuroImage, 124, 32–42. https://doi.org/10.1016/j.neuroimage.2015.08.056

Van Horn, J. D., & Toga, A. W. (2009). Is it time to re-prioritize neuroimaging databases and digital repositories? NeuroImage, 47(4), 1720–1734. https://doi.org/10.1016/j.neuroimage.2009.03.086

White, S. R., Amarante, L. M., Kravitz, A. V., & Laubach, M. (2019). The Future Is Open: Open-Source Tools for Behavioral Neuroscience Research. ENeuro, 6(4), ENEURO.0223-19.2019. https://doi.org/10.1523/ENEURO.0223-19.2019

